# Environmental and morphological constraints interact to drive the evolution of communication signals in frogs

**DOI:** 10.1101/2020.04.18.047936

**Authors:** Matías I. Muñoz, Sandra Goutte, Jacintha Ellers, Wouter Halfwerk

## Abstract

Animals show a rich diversity of signals and displays. Among the many selective forces driving the evolution of communication between individuals, one widely recognized factor is the structure of the environment in which signals are produced, transmitted and received. In particular, animals communicating by sounds often emit acoustic signals from specific locations, such as high up in the air, from the ground or in the water. The properties of these different display sites will impose different constraints on sound production and transmission and may therefore drive signal evolution. Here, we used comparative phylogenetic analyses to assess the relationship between the display site properties and the structure of reproductive calls from 161 frog species from the frog families Ranidae, Leptodactylidae and Hylidae. Specifically, we compared the dominant frequency of species that vocalize from aquatic versus non-aquatic sites, and its relation with body size. We found that the dominant frequency of frogs calling from the water was lower than that of species calling outside of the water, a trend that was consistent across the three families studied. Furthermore, phylogenetic path analysis revealed that the call site had both direct and indirect effects on the dominant frequency. Indirect effects were mediated by call site influencing male body size, which in turn was negatively associated to call dominant frequency. Our results suggest that properties of display sites can drive signal evolution, most likely through morphological constraints, in particular the ones imposed on the sound production mechanism. Also, variation in body size between calling sites explained some of the differences we found in call frequency, highlighting the relevance of the interplay between morphological adaptation and signal evolution. Changes of display site may therefore have important evolutionary consequences, as it may influence sexual selection processes and ultimately may even promote speciation.

**Impact summary:** To attract or impress mates, animals have evolved a great diversity of communication signals, such as song and dance, or brightly colored body parts. Whether these sexual signals are successful depends to a large extent on the environment in which they are produced, transmitted and perceived. For acoustic signals, such as the mating calls of frogs, the environment is well known to influence both their transmission and perception. The impact of the environment on the production of sounds is however far less understood. Here we studied the relation between the environment and signal design across a wide range of frog species, specifically comparing calls of aquatic versus non-aquatic species.

Frogs that called from water were found to call at lower pitch, which was partly explained by the fact that they were also larger. Our results point towards an important environmental driver of signal evolution, namely morphological constraints on signal production. We argue that the environment can impose limits on morphological traits that are either directly or indirectly involved in signal production. Such a mechanism would in particular be important when species move into new habitats, as rapid changes to display sites may lead to rapid changes in sexual signaling and sexual attractiveness.

## Introduction

Animals communicate with an extraordinary variety of display behaviors that span most sensory modalities (Bradbury & Vehrencamp 2011; Stevens 2013). These chemical, visual or acoustic signals are known to experience strong selection pressures imposed by intended and unintended receivers, in particular in the context of sexual communication. The balance between sexual (e.g. mates) and natural selection pressures (e.g. eavesdropping predators) is, however, not independent from the environment. By displaying from sites with particular properties, such as locations with reduced exposure to predators, animals can alter the selection pressures operating on their signals, and thus their evolution.

Irrespective of the sensory modality, one common feature of communication systems is the presence of three interacting components: a sender that produces a signal, a receiver that perceives it, and the transmission environment in between them (Bradbury & Vehrencamp 2011). Environmentally-dependent selection pressures can operate in any of these three processes, with important consequences for signal evolution. In the case of acoustic signals, several studies have investigated the role of variation in transmission environment as a driving factor of signal evolution (e.g., Richards & Wiley 1980; Peters & Peters 2010; García-Navas & Blumstein 2016; Derryberry *et al.* 2018). During transmission, sound signals will experience changes in their temporal and spectral properties (Bradbury & Vehrencamp 2011), which can affect the capacity of receivers to process these signals. Efforts to link acoustic signal features to optimal transmission properties, however, have not yielded consistent results (Ey & Fischer 2009).

From the receiver’s perspective, the presence of noise is another environmental factor relevant for signals evolution. The capacity of receivers to detect and process a signal will be compromised by the presence of background noise. Thus, spectral overlap between noise (e.g., the sounds of other organisms, stream noise or anthropogenic noise) and animals signals is thought to drive changes in the frequency content of sounds produced by senders (e.g., Ryan & Brenowitz 1985; Slabbekoorn 2004; Brumm & Slabbekoorn 2005; Halfwerk *et al.* 2011; Goutte *et al.* 2016). While the transmission environment and background noise can influence signal evolution because they affect the perception of receivers, whether the environment can have more direct effects on the sound production mechanisms is far less understood.

Perhaps, because the morphology of the sound producing structures is generally considered to impose strong constraints, direct environmental influences on sound production have received less attention, in particular in the case of vocalizations. Still, acoustic signal production can be influenced by external factors. The environment can influence the biomechanics of sound production through changes in a sender’s physiology. In ectotherms, the temporal structure of acoustic signals is strongly determined by the environmental temperature (Cusano *et al.* 2016; Ziegler *et al.* 2016). Alternatively, the environment immediately surrounding a signaler imposes constraints on the biomechanics of sound-producing organs. The production of vocalizations generally involves changes in body posture and the inflation/deflation of body parts, and environmental constraints on any of these processes will also impact the signal (Halfwerk *et al.* 2017). Interestingly, some animals can manipulate their environment to release them from the constraints imposed by their morphology. In tree-crickets the wings are too small relative to the wavelength of the sounds they produce, resulting in poor sound radiation. By modifying leaves to act like acoustic baffles and using them as calling site these insects overcome the morphological constraint on signal production, greatly improving sound radiation (Mhatre *et al.* 2017).

Anurans (frogs and toads) represent an excellent group to study the influence of the environment on signal production because of the diversity of calling sites that different species use. Males advertise their readiness to mate to females by calling from the water, while floating, sitting or being submerged, or from land, while sitting on rocks, vegetation or in burrows (Wells 2007). Furthermore, the acoustic properties of advertisement calls are species-specific and closely linked to the biomechanics of sound production. The spectral content of calls will be determined by the morphology of the larynx (e.g., Baugh *et al.* 2018; López *et al.* 2020) and the pattern of vocal sac inflation (e.g., Dudley & Rand 1991; Zhang *et al.* 2016). Different calling sites occupied by frogs will impose different constraints on sound production. For example, frogs calling from shallow water cannot inflate their lungs and vocal sacs to the same extent as freely floating individuals, a biomechanical constraint accompanied by a number of changes in the calls, including the production of higher frequency vocalizations (Goutte *et al.* 2020). These signal modifications caused by calling site properties have important consequences for mate attraction and, thus have the potential to drive signal evolution (Halfwerk *et al.* 2017).

In the present study we used comparative phylogenetic methods to study the effects of calling site on the evolution of frog vocalizations. We compared the dominant frequency of species that call from inside and outside of the water, and evaluated how these variables relate to body size. We hypothesized that aquatic calling sites will impose fewer mechanical constraints on the production of calls than non-aquatic sites, and thus we expected higher frequency calls in species using water to vocalize. This hypothesis is supported by previous intra-specific experiments on frogs calling from deep and shallow water (e.g., Goutte *et al.* 2020), but has not been evaluated in a comparative framework.

## Methods

### Data collection and categorization

We restricted the data collection to species from the families Ranidae, Leptodactylidae and Hylidae present in the molecular phylogeny published by Pyron (2014). We chose these families because they are species-rich clades relative to other frog families (more than 200 species in each family, Frost 2020), span wide geographic distributions, and are known to occupy both aquatic and non-aquatic calling sites. These families are not closely related to each other (not sister clades), and include species with diverse lifestyles and ecomorphologies. Also, the vocal behavior of species in these families has been investigated with some detail, and data on call frequency and body size are available from the literature.

For each family, we collected data on the snout-vent length, dominant frequency and calling site. Most of the information was obtained from the literature or other digital sources (see below). Personal measurements made by the authors of the present article were also included. If searched in the literature, body size and call dominant frequency were obtained from other comparatives studies and books. We restricted our search to body size of males and the dominant frequency of advertisement vocalizations. Information on calling sites was obtained mainly from verbal descriptions of frog vocal behavior present the literature, the specialized website AmphibiaWeb, and from the personal experience of the authors. Multimedia information available from AmphibiaWeb and Youtube, such as pictures and videos of calling males, was used to confirm ambiguous verbal descriptions. For a few species, multimedia information was used as the sole criterion for calling site assignment. Each species was assigned to one of three possible calling site categories: (1) aquatic, (2) non-aquatic, and (3) mixed. Aquatic species included frogs that vocalize either standing in water, or floating on the water surface. The non-aquatic category included species that call from the ground, or from perched positions on trees or rocks without direct contact with water. Species calling from cavities dug in the ground or cavity-like structures on vegetation (e.g., the axils of bromeliads) were also included in the non-aquatic category. The few species for which both aquatic and non-aquatic calling was described were assigned to the mixed category. In total, we collected body size, dominant frequency and calling site data for 51 Ranidae, 54 Leptodactylidae, and 71 Hylidae species. Phylogenetic trees of each family showing the data for each species are shown in Figure 1.

**Figure 1:**
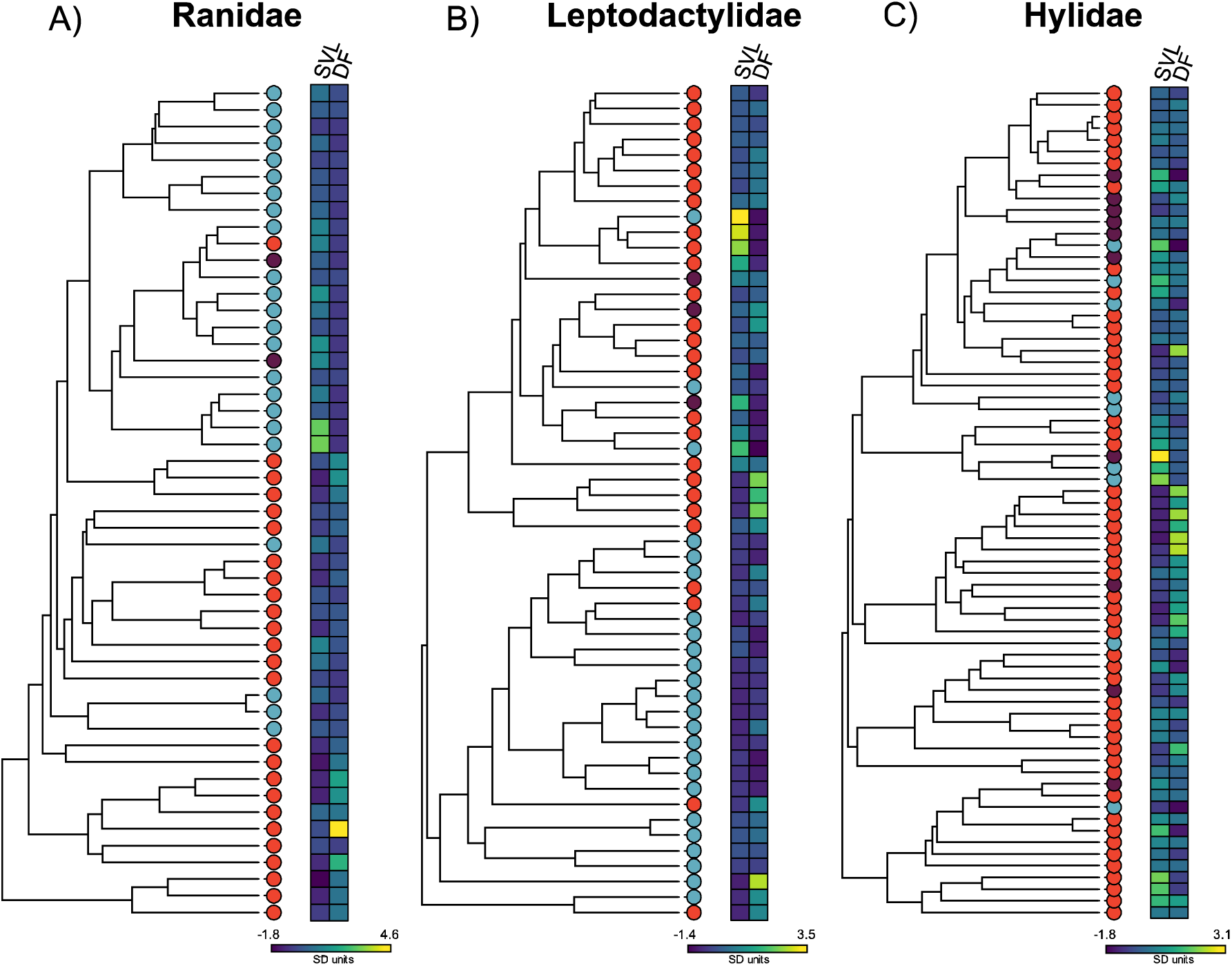
Phylogenetic trees of A) Ranidae, B) Leptodactylidae, and C) Hylidae. Colored circles next to the tips of the trees depict species that call from the water (blue), out of the water (red) or in the mixed category (purple). Body size (SVL) and dominant frequency (DF) data are plotted next to each tree. SVL and DF values were transformed to standardized SD units for visualization purposes.

### Comparative analyses

All the analyses were performed in R (version 3.6.1, R Core Team 2019). We pruned the phylogenetic tree of Pyron (2014) to exclude all the species that were not present in our data set. Before performing the analyses, we excluded the species in the mixed calling site category (Ranidae N = 2, Leptodactylidae N = 3, and Hylidae N = 10). These corresponded to a small subset of the species, and were excluded because one of the analyses allows only binary categorical variables, and because we were mainly interested in the aquatic versus non-aquatic comparison. We used phylogenetic generalized least squares (PGLS) to evaluate the effect of call site and body size on call dominant frequency. For each frog family we fitted a separate PGLS model, and all the models included the log_10_-transformed dominant frequency as response variable, and the log_10_-transformed body size and calling site (‘aquatic’ vs ‘non-aquatic’) as explanatory variables. The interaction between calling site and body size was not significant for any family, and we removed it from the models before computing the coefficients reported here. We used the library ‘*ape*’ (version 5.3, (Paradis & Schliep 2019) to create a correlation structure assuming a Brownian motion model of trait evolution, which was then used to fit the PGLS models in R. Plots of residuals versus fitted values and residual quantile-quantile were used to evaluate departures from regression assumptions.

To further explore the causal relationships between dominant frequency, body size and calling site we used phylogenetic path analysis (PPA, von Hardenberg & Gonzalez-Voyer 2013) implemented in the library ‘*phylopath*’ (version 1.1.1, van der Bijl 2018). We *a priori* defined three hypotheses describing the causal relationships between these variables (Fig. 2). The first hypothesis included only a direct path linking body size and call dominant frequency (Fig. 2a). We consider this our null model because it excludes any influence of calling site on body size or on dominant frequency. This path was retained in the other alternative models because the negative association between call frequency and body size is a well-described pattern in animal vocal sound production. Hypothesis 2 and 3 included a direct path linking calling site and body size (Fig. 2b), and a direct link between calling site and dominant frequency (Fig. 2c), respectively. For each family the three models were compared based on their CICc information criterion value. In case more than one model was best ranked (i.e., more than one model within ΔCICc < 2 from the top ranked model), we used conditional model averaging (i.e., missing paths are not included in the average) to obtain a single average model. Similar to the PGLS analyses, we assumed a Brownian motion model of trait evolution for the linear models underlying the PPA analyses. Additionally, we also performed the path analysis after pooling together the data collected for the three families into a single data set and phylogenetic tree containing the N=161 species. For this analysis we tested the same set of models (Fig. 2), and followed the same procedure used for the analyses of the three families by separate.

**Figure 2:**
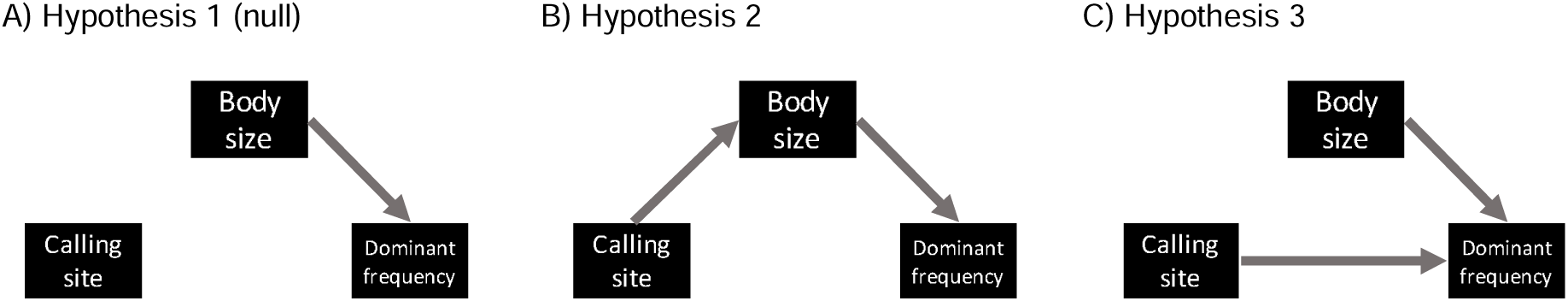
Directed acyclic graphs (DAG) describing the three causal hypotheses tested using phylogenetic path analysis.

## Results

### Phylogenetic Generalized Least Squares Analysis (PGLS)

Male body size and call dominant frequency were negatively associated in the three families studied (Table 1). For the families Leptodactylidae and Hylidae, the frogs that called from the water did so at lower dominant frequencies than non-aquatic species (Table 1, Fig. 3d, c). A similar trend was followed by Ranidae species, though differences between aquatic and non-aquatic frogs were not significant (Table 1, Fig. 3a).

**Table 1:** Results of PGLS models fitted for the three families. In all the models the dependent variable was log_10_-transformed dominant frequency. Bold numbers depict significant results.

| Family | Variable | Estimate | S.E. | t-value | P-value |
| --- | --- | --- | --- | --- | --- |
| Ranidae |  |  |  |  |  |
|  | Intercept | **5.57** | **0.45** | **12.36** | **< 0.0001** |
| | $\log_{10}(\text{SVL})$ | **-1.31** | **0.25** | **-5.28** | **< 0.0001** |
|  | Display site: *Non-aquatic* | 0.11 | 0.11 | 0.96 | 0.3418 |
| Leptodactylidae |  |  |  |  |  |
|  | Intercept | **4.60** | **0.30** | **15.15** | **< 0.0001** |
| | $\log_{10}(\text{SVL})$ | **-0.97** | **0.19** | **-5.02** | **< 0.0001** |
|  | Display site: *Non-aquatic* | **0.23** | **0.07** | **3.37** | **0.0015** |
| Hylidae |  |  |  |  |  |
|  | Intercept | **4.04** | **0.39** | **10.3** | **< 0.0001** |
| | $\log_{10}(\text{SVL})$ | **-0.57** | **0.23** | **-2.45** | **0.0175** |
|  | Display site: *Non-aquatic* | **0.26** | **0.09** | **2.96** | **0.0044** |

**Figure 3:**
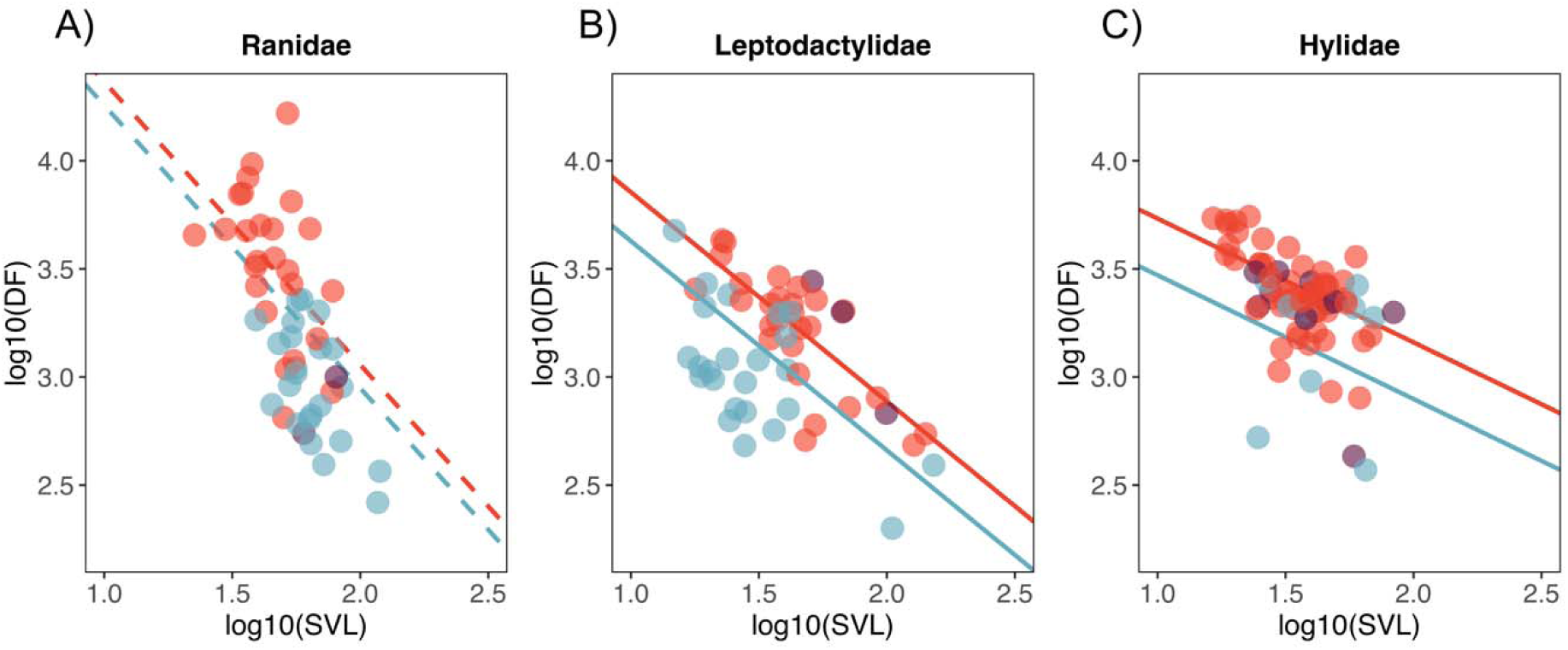
Scatterplot showing the association between body size, dominant frequency, and the effect of calling site for A) Ranidae, B) Leptodactylidae and C) Hylidae. Colors blue, red and purple correspond to species in the aquatic, non-aquatic, and mixed calling site categories. Points represent the raw data, and regression lines represent PGLS model estimates. Species in the mixed calling site category are shown but were not included in the PGLS analyses. The dashed lines in Ranidae depict non-significant differences between the intercepts of aquatic and non-aquatic frogs.

### Phylogenetic path analysis

When families were analyzed separately, the causal hypotheses that include a direct effect of calling site on body size (Hypothesis 2), and on dominant frequency (Hypothesis 3) were among the best ranked models (i.e., within ΔCICc < 2) (Table 2). For Ranidae, the null hypothesis where there is no effect of display site on dominant frequency or on body size was also supported (Table 2). Model averaging of the best ranked models showed that the three families followed the same general pattern of causal association between the variables (Fig. 4a). That is, calling site had a positive and direct effect on dominant frequency, and a negative direct effect on body size. Nevertheless, the coefficient estimates for these paths did not significantly deviate from zero, as judged by the 95% confidence intervals overlapping with 0 (Fig. 4a). As expected, body size had a negative effect on dominant frequency, and was the only significant path for the three families (Fig.4a).

**Table 2:** Summary of best ranked models tested using phylogenetic path analysis. The set of models evaluated is shown in Fig. 2. Bold numbers depict the best set of causal models (within ΔCICc < 2) that were latter used for model averaging. C = Fisher’s C statistic; k = number of independence claims; q = number of parameters; ΔCICc = difference in CICc from the top ranked model; CICc weights = model conditional weight.

| Family | Model | k | q | C | P-value | CICc | $\Delta\text{CICc}$ | Likelihood | CICc weights |
| --- | --- | --- | --- | --- | --- | --- | --- | --- | --- |
| Ranidae |  |  |  |  |  |  |  |  |  |
|  | **H3** | **1** | **5** | **1.25** | **0.535** | **12.7** | **0.00** | **1.00** | **0.474** |
|  | **H1** | **2** | **4** | **4.40** | **0.355** | **13.3** | **0.65** | **0.724** | **0.343** |
|  | **H2** | **1** | **5** | **3.15** | **0.207** | **14.6** | **1.89** | **0.388** | **0.184** |
| Leptodactylidae |  |  |  |  |  |  |  |  |  |
|  | **H2** | **1** | **5** | **3.17** | **0.205** | **14.5** | **0.00** | **1.00** | **0.594** |
|  | **H3** | **1** | **5** | **4.99** | **0.083** | **16.3** | **1.82** | **0.40** | **0.239** |
|  | H1 | 2 | 4 | 8.16 | 0.086 | 17.0 | 2.53 | 0.28 | 0.168 |
| Hylidae |  |  |  |  |  |  |  |  |  |
|  | **H3** | **1** | **5** | **3.99** | **0.136** | **15.1** | **0.00** | **1.00** | **0.526** |
|  | **H2** | **1** | **5** | **4.94** | **0.085** | **16.0** | **0.95** | **0.62** | **0.328** |
|  | H1 | 2 | 4 | 8.92 | 0.063 | 17.6 | 2.56 | 0.28 | 0.146 |
| All families |  |  |  |  |  |  |  |  |  |
|  | **H3** | **1** | **5** | **6.90** | **0.032** | **17.3** | **0.00** | **1.00** | **0.507** |
|  | **H2** | **1** | **5** | **7.14** | **0.028** | **17.5** | **0.24** | **0.89** | **0.451** |
|  | H1 | 2 | 4 | 14.04 | 0.007 | 22.3 | 5.01 | 0.08 | 0.042 |

**Figure 4:**
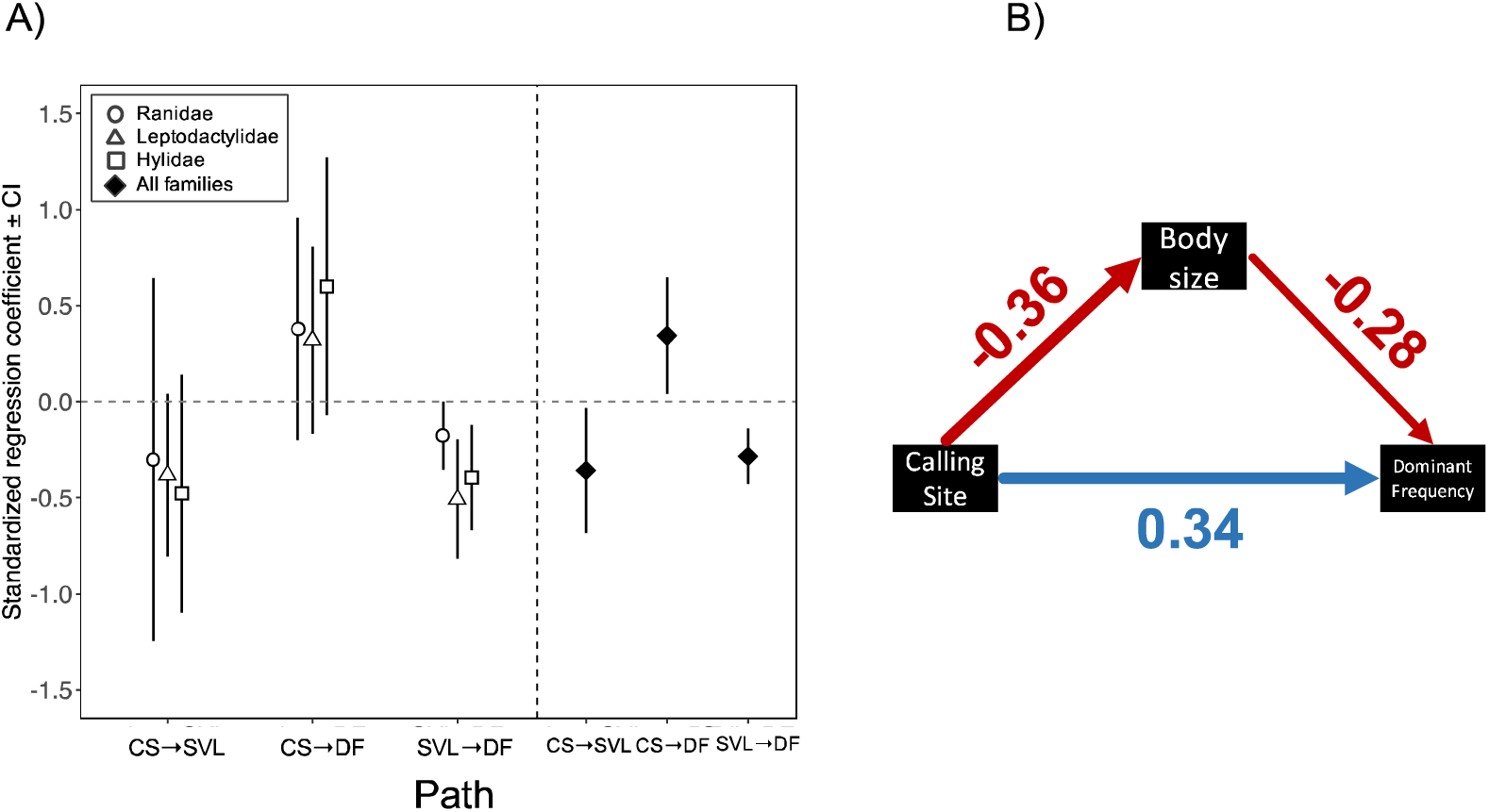
A) Standardized regression coefficients estimated from the average model for each family (open symbols), and for all the families together (filled symbols). Error bars correspond to 95% confidence intervals obtained after 500 bootstrap replications. B) Average model for the three families analyzed together. Numbers correspond to standardized regression coefficients, and are the same as the filled symbols in Fig. 4a. Arrows widths and colors depict the size and the direction of the effect.

The analysis of the three families together revealed the same trend observed for each family separate. The models that included a causal link between calling site and body size (Hypothesis 2), and between calling site and dominant frequency (Hypothesis 3) outweighed a model where calling site was independent of these variables (Hypothesis 1) (Table 2). The average model included a positive and direct effect of calling site on dominant frequency, and a negative direct effect on body size (Fig. 4a, b). It also included a negative effect of calling site on body size, which indicated that calling site can indirectly affect dominant frequency through effects on body size (Fig. 4a, b). In the average model, the estimated coefficients of all the paths were of similar magnitude (range: [0.28-0.34]) and significantly different from zero (Fig. 4a).

## Discussion

Signal evolution is driven by a number of factors, including the environment and the morphology of the sender. Here, we evaluated the impact of calling site on the evolution of call frequency in three frog families using comparative phylogenetic methods. We found that species vocalizing from the water call at lower dominant frequencies than species calling from non-aquatic locations. Furthermore, our analyses revealed that calling site had both direct and indirect effects on call frequency. Because body size and signal frequency were negatively associated in the three families studied, direct effects of calling site on body size had an indirect impact on signal frequency. These results indicate that environmental constraints interact with morphological constraints to drive the evolution of frog vocalizations.

Our comparison across families revealed a similar pattern of call frequency variation as found within species. In the hylid frog *Boana atlantica* the dominant frequency of calls emitted by individuals in the water is, for example, lower than calls produced from vegetation (Camurugi *et al.* 2015). Similarly, floating túngara frogs call at lower dominant frequencies when compared to trials in which individuals are experimentally forced to call while resting on a solid substrate (Goutte *et al.* 2020). Shallow water conditions prevented males from fully inflating their vocal sac in this study, indicating that call site-induced constraints on sound production have an immediate impact on signal frequency (Goutte *et al.* 2020). We extend this argument here, and propose that the frequency differences we found in our analyses are mainly caused by different constraints on sound production imposed by aquatic and non-aquatic calling sites. Furthermore, calling sites differ not only in the biomechanical constraints they impose, but also in other factors like exposure to desiccation or temperature (Camurugi *et al.* 2015; Cicchino *et al.* 2020). This suggests that other call variables, such as some temperature-dependent temporal patterns may also be impacted by calling site choice. Variation in call sites can therefore influence signal production by altering sender’s physiology or biomechanical constraints.

The size of the sound-producing organ can determine the frequency content of vocalizations, and thus larger animals produce lower frequency sounds (Fletcher 2004). The association between size and frequency is a well-described physical consequence of vocal sound production, and it is suggested that ecological factors driving changes in body size also have concomitant effects on signal frequency (Wilkins *et al.* 2013). A similar case of morphology-driven signal evolution can be found in Darwin finches, where diet-dependent changes in beak morphology are accompanied by modification in song production (Podos 2001; Podos & Nowicki 2004). Our analyses show that body size variation in frogs is to some extent explained by the different calling sites they occupy, with consequences for call frequency. Body size evolution in ectotherms has been linked to a number of environmental factors, including temperature, humidity, and evapotranspiration potential (e.g., Amado *et al.* 2019; Velasco *et al.* 2020) all of which are known to differ between calling sites. Furthermore, morphological adaptations are also expected to differ between calling sites. Arboreal or fossorial habits, for example, are linked to a number of morphological specializations (Moen *et al.* 2013), including differences in body size (e.g., Dugo-Cota *et al.* 2019). Many frog species vocalizing out of the water call while sitting on vegetation, perched on branches or leaves. Large frogs may be unable of arboreal calling due to the lack of physical support provided by hanging leaves and branches. In contrast, body size may be less constrained in terrestrial or aquatic calling species. In our data arboreal calling species were included into the non-aquatic calling site category, and were not analyzed separately because they were mostly present in the family Hylidae, but scarce in Ranidae and Leptodactylidae. Still, we predict arboreal species to have even higher frequency calls relative to aquatic and terrestrial species due to a combination of body size constraints, and favorable transmission of high frequencies from elevated sites (Mathevon *et al.* 1996; Schwartz *et al.* 2016; Cicchino *et al.* 2020). Variation in body size will not only have a direct impact on call frequency due to allometry, but may also limit the possible calling sites a species can occupy, highlighting the relevance of the interaction between morphology and calling site on signal evolution.

Display sites can drive signal evolution through direct impacts on the sound production mechanism, as well as other selection pressures on senders and receivers. Divergent transmission properties between display sites may select for signals with matching properties over evolutionary time scales. Likewise, signal adaptation to display site-dependent noise profiles may also operate on the long-term. Environmental constraints on production mechanisms will however have immediate consequences when senders move to different display sites. For example, a frog that moves from an aquatic to a non-aquatic calling site will immediately call at higher frequencies. Therefore, the impact of the environment, and in particular of display sites, on vocal production mechanisms has the potential to cause fast signal divergence, and in some cases promote speciation.

## Acknowledgements

MIM is supported by Becas Chile 2018-CONICYT scholarship (number: 72190501). The authors declare no conflicts of interest.

## Author contributions

SG, JE and WH conceived and designed the study. MIM, SG and WH collected the data. MIM and SG analyzed the data. MIM and WH drafted the initial version of the manuscript and all the authors commented and edited later versions of the manuscript.

## Data accessibility

Data will be made available on Dryad upon acceptance.

